# Non-obese genetic type 2 diabetes causes brain and behavioral hallmarks of chronic stress

**DOI:** 10.1101/2025.06.08.658427

**Authors:** Marina Romaní-Pérez, Mathilde S. Henry, Clémentine Pajot, Danielle Bailbé, Julie Brossaud, Khulganaa Buyannemekh, Lucia Prado, Aline Foury, Marion Rincel, Amandine L. Lépinay, Bernard Portha, Jonathan D. Turner, Jamileh Movassat, Muriel Darnaudéry

**Affiliations:** Univ. Bordeaux, INRAE, Bordeaux INP, NutriNeuro, UMR 1286, F-33000, Bordeaux, France; Univ. Paris Cité, Unit of Functional and Adaptive Biology - CNRS UMR 8251, Team “Biology and Pathology of the Endocrine Pancreas (B2PE); Department of Infection and Immunity, Luxembourg Institute of Health (LIH), Esch-sur-Alzette, Luxembourg

**Keywords:** Goto-kakizaki rats, glucocorticoids, medial prefrontal cortex, adrenalectomy, depression, anxiety

## Abstract

The comorbidity of obesity, type 2 diabetes (T2D), and psychiatric disorders— particularly anxiety and depression—is well documented. However, it remains unclear whether T2D, independently of obesity, contributes to the development of emotional dysfunctions. Furthermore, alterations in the hypothalamic-pituitary-adrenal (HPA) stress axis are commonly associated with both T2D and depression, but the role of stress in emotional disorders linked to T2D has been poorly explored. This study aimed to investigate the impact of T2D, independent of obesity, on the neuroendocrine stress axis, as well as molecular, cellular, and behavioral indicators of emotional dysfunction. Using the non-obese Goto-Kakizaki (GK) rat model of T2D, we assessed the effects of diabetes on hormonal and neuronal stress responses, molecular and structural markers of stress in the brain, and anxiety- and depressive-like behaviors. We also evaluated the impact of adrenalectomy in GK rats to determine the contribution of glucocorticoids to their behavioral impairments. Our findings reveal that non-obese diabetes leads to heightened endocrine and brain responses to stress, along with upregulation of stress-related molecular markers and structural features indicative of chronic stress, particularly in the medial prefrontal cortex. Additionally, GK rats exhibited pronounced anxiety- and depressive-like behaviors. Importantly, lowering glucocorticoid levels in GK rats helped alleviate some of the metabolic and emotional disturbances. This study suggests that T2D, independent of obesity, induces stress-related brain and behavioral changes, partly mediated by glucocorticoids.

**Highlights:**

- T2D disrupts endocrine and neural responses to stress.
- T2D alters stress-related gene expression in the medial prefrontal cortex.
- T2D induces structural changes in the medial prefrontal cortex.
- T2D contributes to anxiety- and depressive-like behaviors.
- Reducing corticosterone levels mitigates anxiety-like behavior in diabetic rats.

## 1. Introduction

According to the 2021 Global Burden of Disease report, the leading causes of disability worldwide include depression, anxiety disorders, and diabetes [1]. More than 90% of diabetic patients are diagnosed with type 2 diabetes (T2D), primarily in adults, but its incidence is also rising among younger populations. The consequences of T2D on the brain in these younger populations, however, are not yet fully understood. The major hallmark of T2D is elevated blood glucose and insulin resistance [2], but it is also frequently associated with other metabolic disturbances such as dyslipidemia, hypertension and obesity [3]. Depression and anxiety disorders are more prevalent among diabetic patients compared to the general population, and these emotional disorders are also associated with increased risk for T2D [4]. Growing evidence suggests shared pathophysiological mechanisms between T2D and anxiety or depressive disorders, including inflammation and hyperactivity of the hypothalamic-pituitary-adrenal (HPA) axis [4,5]. T2D is increasingly diagnosed at younger ages [6,7] and in these cases, it is not necessary associated with obesity [6,8]. However, the obesity-independent impact of T2D on brain and neuropsychiatric symptoms is not yet fully understood.

In preclinical studies, a substantial body of evidence suggests that T2D affects the brain, leading to cognitive and emotional disturbances, particularly exacerbating anxiety and/or depression-like behaviors [9–14]. However, most studies have used overweight T2D models, which involve long-term exposure to high-fat or high-sugar diets [10,12,15–17] or mutations in leptin or leptin receptors [13]. Given that not all T2D patients are overweight [18] it is essential to explore lean diabetic animal models to better understand the link between T2D and emotional disorders. Additionally, there is growing evidence that T2D has a strong genetic component (in addition to key environmental factors), which has been underexplored in the context of neuropsychiatric diseases. The Goto-Kakizaki (GK) diabetic rat line was developed from normoglycemic Wistar rats by repeated inbreeding of the siblings with the highest blood glucose levels during an oral glucose tolerance test [19]. GK rats are lean and develop a stable, spontaneous T2D phenotype after weaning, characterized by mild hyperglycemia, glucose intolerance, defective insulin secretion, peripheral insulin resistance, and peripheral tissue inflammation [20]. This model offers a valuable tool to study brain alterations and emotional dysfunctions associated with T2D, while eliminating obesity and nutritional confounding factors. Previous research has demonstrated cognitive impairments in GK rats [21] coupled with neurovascular abnormalities [22–24] but their emotional function has been less extensively explored.

Exposure to chronic stress is a known key risk factor for mental health, particularly in the development of anxiety and mood disorders [25]. Interestingly, both obesity and T2D produce alterations in the HPA axis and brain changes that resemble those observed following chronic stress exposure [16]. Previous studies have shown that glucocorticoid overexposure contributes to cognitive disturbances associated with T2D [26,27], but its role in emotional deficits associated to T2D has not fully examined.

In this study, we examined the impact of T2D in the non-obese GK model on the neuroendocrine stress axis, as well as on molecular and cellular brain features related to stress. We then investigated emotional behaviors in GK rats using a comprehensive set of well-established paradigms for anxiety- and depressive-like behaviors. Finally, we studied emotional functions in diabetic GK rats that had undergone adrenalectomy and received low-dose corticosterone replacement to restore normal basal circadian rhythm.

## 2. Methods and materials

### 2.1 Animals

Male Wistar and GK rats (1.5-3 months old) were housed at 22±2 °C with free access to food (SAFE D113, Augy, France) and water in a 12h light/dark cycle [“lights on”: Zeitgeber Time (ZT)0]. All experimental procedures are in line with the European Union guidelines for the use of animals for experimental purposes (Council Directive 2010/63/EU) and with the French ones (Directive 87/148, Ministère de l’Agriculture et de la Pêche – Apafis #16924).

### 2.2 Glycemia, glucose tolerance test and insulin secretion

Glycemia was measured at ZT3.5 and ZT12.5. At ZT8, an intraperitoneal glucose tolerance test (IPGTT) was conducted to measure glycemia and insulin in plasma before and after intraperitoneal glucose administration. Details are provided in supplementary methods.

### 2.3 Corticosterone measurements

Circadian variations of corticosterone levels were measured in plasma obtained from blood collected from the lateral tail vein at ZT8 and ZT12.5. Blood was also collected to evaluate the HPA-axis function by measuring plasma corticosterone levels at baseline and 10 and 120 min after a mild stress (i.e 10 min in an open-field). Additionally, corticosterone was measured in isolated brain areas [i.e prefrontal cortex (PFC), hypothalamus, amygdala and hippocampus]. Detailed methods are provided in supplementary methods.

### 2.4 real-time qPCR

RNA extraction, reverse transcription and real-time qPCR were performed as explained in supplementary methods.

### 2.5. Golgi-cox staining

Golgi-cox staining was conducted in brains to analyze dendritic arbor complexity and relative spine number (4-6 neurons per animal) of pyramidal neurons in layer II/III of the medial (m)PFC as previously described [28] (more details are provided in supplementary methods).

### 2.6 Immunohistochemistry

The evaluation of astrocytes in the mPFC and activated neurons in mPFC, paraventricular nucleus (PVN), amygdala and hippocampus was performed through immunohistochemistry procedures as described in supplementary methods.

### 2.7. Anxiety-like and depressive like behaviors

Wistar and GK rats’ behavior was studied using a battery of tests relevant for psychiatric disorders: light-dark test, open-field, tunnel test for anxiety-like behaviors, social interaction, forced swimming test (FST) and locomotor circadian rhythm for depressive-like behaviors. Details of the behavioral tests are given in the supplementary methods.

### 2.8. Peripheral glucocorticoids normalization experiment

Bilateral adrenalectomy or sham surgery were conducted in anesthetized GK rats by a dorsal approach. Low-dose corticosterone replacement (25µg/mL in 0,9% saline) was given via the drinking water. Measurements of corticosterone, glycemia, insulin and behavior are detailed in supplementary methods.

### 2.9. Statistical analysis

Nine different batches were used in the study to assess effects of non-obese diabetes on metabolism, corticosterone levels, brain and behavior and the impact of ADX in GK rats. Body weight and basal glycemia were systematically measured in all cohorts. Sample sizes were determined based on power analysis and common practice in behavioral (∼8-10 animals per group) and cellular and molecular biology (∼5 animals per group) experiments. Statistical outliers were detected with the Grubb’s test and highly significant outliers (P<0.01) were removed from analyses. For the ADX experiment, 3 animals were excluded from the study because they were not completely adrenalectomized (as seen by high corticosterone levels post stress). Data quantifications that potentially include subjective bias (social interaction, light/dark, tunnel test, EGR-1, GFAP, Golgi quantification) were conducted by observers blind to the experimental group. Data were analyzed with Statistica (Statsoft, TULSA, OK, USA) or GraphPad version 7.0 (La Jolla, CA, USA). Normality was assessed using Shapiro–Wilk tests. Unpaired Student’s *t*-tests were conducted for comparisons between 2 independent groups. Two-way ANOVA with repeated measures was used to assess interactions between time and genotype and main effects. Tukey’s post-hoc test or planned comparisons were performed when interactions were identified. Statistical significance was set at p<0.05.

## 3. Results

### 3.1 Goto-Kakizaki rats exhibit type 2 diabetes phenotype but are not overweighed

As expected, GK rats showed hyperglycemia compared with Wistar controls both during light (ZT3.5; t_(19)_=6.23, p<0.0001) and dark phase (ZT12.5; t_(13)_=9.25, p<0.0001) (**Figure 1A**). GK rats showed lower body weight (t_(32)_=11.59, p<0.0001; **Figure 1B**) and size (t_(16)_=5.73, p<0.0001; **Figure1C**) than age-matched Wistar rats. In response to intraperitoneal glucose administration during the IPGTT, a time and a genotype effect were identified for glycemia (two-way ANOVA with repeated measures; Time effect: F_(5, 80)_=44.77, p<0.0001; Genotype effect: F_(1,16)_=61.94, p<0.0001; **Figure 1D**), as well as a significant genotype x time effect (F_(5,80)_=26.93, p<0.0001; **Figure 1D**). Contrary to Wistar rats, that showed a slight increase of their glycemia at 15 min (Tukey’s post-hoc, p=0.0085 versus T0) returning to the basal levels at 60 min (p=0.06 versus T0), GK rats had a marked glycemia increase 15 min after glucose administration (p<0.0001 versus T0), remaining elevated 120 min post-administration (p=0.018 versus T0) (**Figure 1D**). Accordingly, the area under the curve of the IPGTT was higher in GK than in Wistar rats (t_(10)_=10.79, p<0.001, **Figure 1D**, insert) indicating glucose intolerance. Insulinemia was also higher in fasted GK rats (basal) (t_(15)_=3.89, p<0.01; **Figure 1E**) and differentially regulated between genotypes during the IPGTT (genotype x time, F_(3,45)_=18.02, p<0.0001; **Figure 1E**). While glucose administration efficiently increased insulin secretion in Wistar rats (Tukey’s post-hoc p<0.0001, T15 versus T0), insulin levels remained unchanged in GK rats (p=0.998 at 15 min) (**Figure 1E**). Overall, these data validate the use of GK rats for the study of T2D-related metabolic disturbances without the confounding effects of obesity.

**Figure 1.**
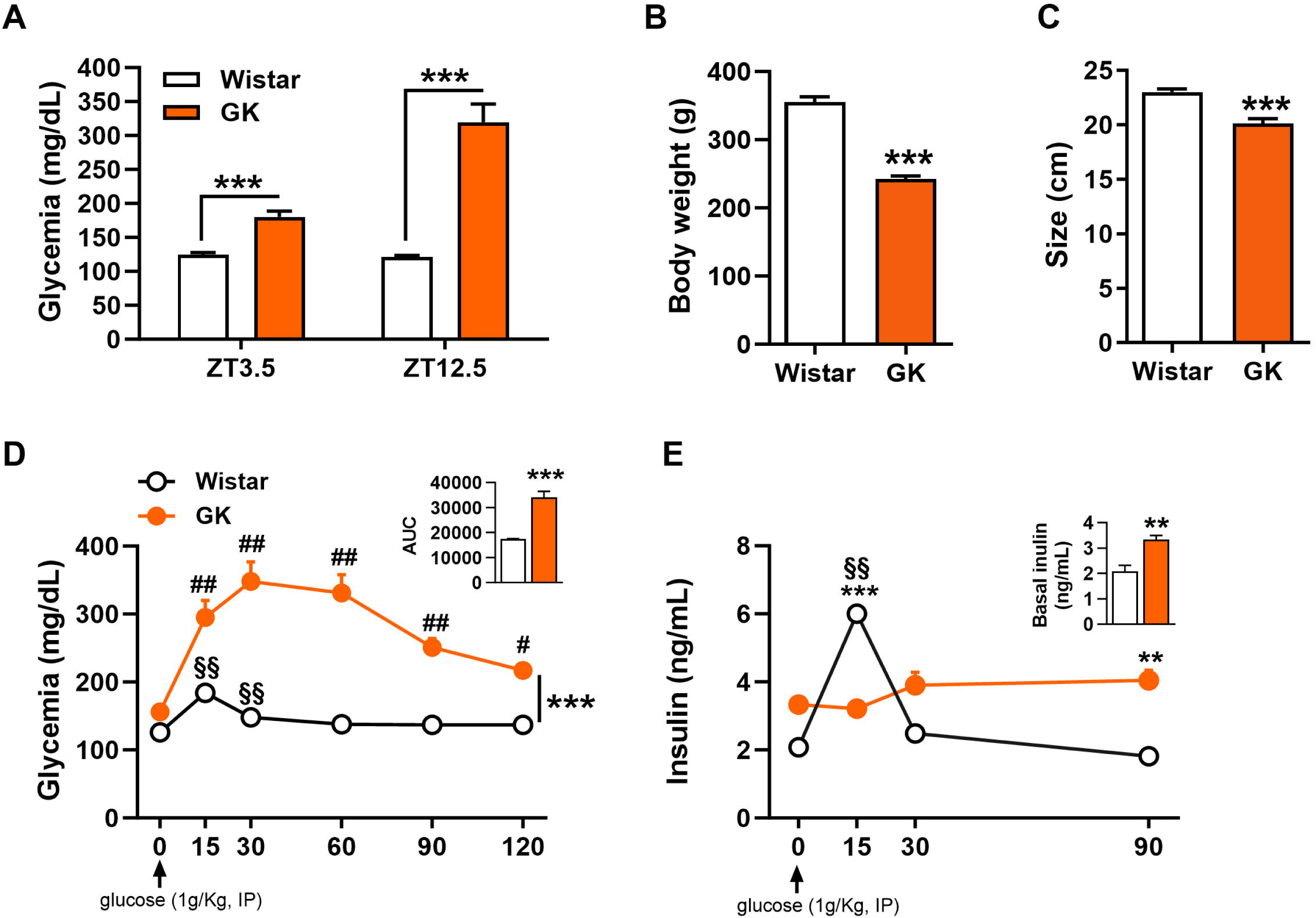
Goto-kakizaki (GK) rats show type 2 diabetes (T2D) without obesity. **(A)** GK rats exhibited non-fasted hyperglycemia both at zeitgeber time (ZT) 3.5 (Wistar, n=18; GK n=16) and ZT12.5 (Wistar n=9; GK, n=6). ZT0 refers to lights on and ZT12 to lights off. **(B)** GK rats had reduced body weight (Wistar, n=18; GK n=16) and **(C)** small size than Wistar rats (Wistar, n=8; GK n=10). **(D)** Impaired whole-body glucose clearance in GK rats after an intraperitoneal (IP) administration of glucose in the glucose tolerance test (GTT) with higher area under the curve (AUC) (Wistar, n=10; GK n=8). **(E)** Increased basal insulin and reduced insulin secretion in response to glucose (IP) in GK rats (Wistar, n=10; GK n=7). Data are expressed as mean ± SEM. Unpaired Student’s t-test: (A-C), AUC and basal insulin in (C) and (E); Two-way ANOVA with repeated measures followed by Tukeýs post-hoc test: (D and E). **p<0.01 and ***p<0.001 vs Wistar rats; ^§§^p<0.01, each time point vs T0 in Wistar rats; ^#^p<0.05 and ^##^p<0.01, each time point vs T0 in GK rats.

### 3.2 Goto-kakizaki rats display a hyperactive hypothalamic-pituitary-adrenal stress axis, elevated brain corticosterone levels and changes in neuronal activation after stress

Based on the critical role of the HPA axis dysfunctions in metabolism and diabetes, we examined glucocorticoids levels in basal and stressed conditions in Wistar and GK rats. Plasma corticosterone levels differently varied throughout ZT between GK and control Wistar rats (two-way ANOVA with repeated measures: genotype x ZT effect, F_(1,12)_=5.205, p<0.05; **Figure 2A**). GK rats exhibited higher plasma corticosterone levels than Wistar control during the inactive light phase (Tukeýs post-hoc, ZT8 p<0.001), whereas groups showed similar plasma corticosterone levels during the dark phase (active phase, ZT12.5) (**Figure 2A**). We then investigated the neuroendocrine response to a mild stress (10 min novelty exposure). Relative to Wistar rats, GK rats showed a higher activation of HPA axis in response to a mild stress (**Figure 2B**; genotype effect, F_(1,14)_=46.86, p<0.0001; time effect, F_(2, 25)_=82.93, p<0.0001 and genotype x time effect, F_(2,25)_=8.58, p<0.01). Post-hoc tests revealed that GK rats had higher plasma corticosterone levels than Wistar rats 10 (p<0.0001) and 120 min (p<0.001) after the stress exposure. Basal plasma corticosterone levels (T0) were also elevated in GK relative to Wistar controls (t_(11)_=5.80, p<0.0001; **Figure 2B**). Whatever the brain area considered, brain corticosterone levels were higher in GK than Wistar rats (PFC, t_(19)_=2.30, p<0.05, hypothalamus, t_(17)_=2.56, p<0.05, amygdala, t_(17)_=2.95, p<0.01 and hippocampus, t_(17)_=2.94, p<0.01, **Figure 2C**). These results suggest that T2D independently of obesity may produce a hyperactivity of the HPA axis in basal and stress conditions. More importantly, our results suggest that the brains of subjects with T2D are exposed to glucocorticoid levels which are twice as high as those of control Wistar subjects.

**Figure 2.**
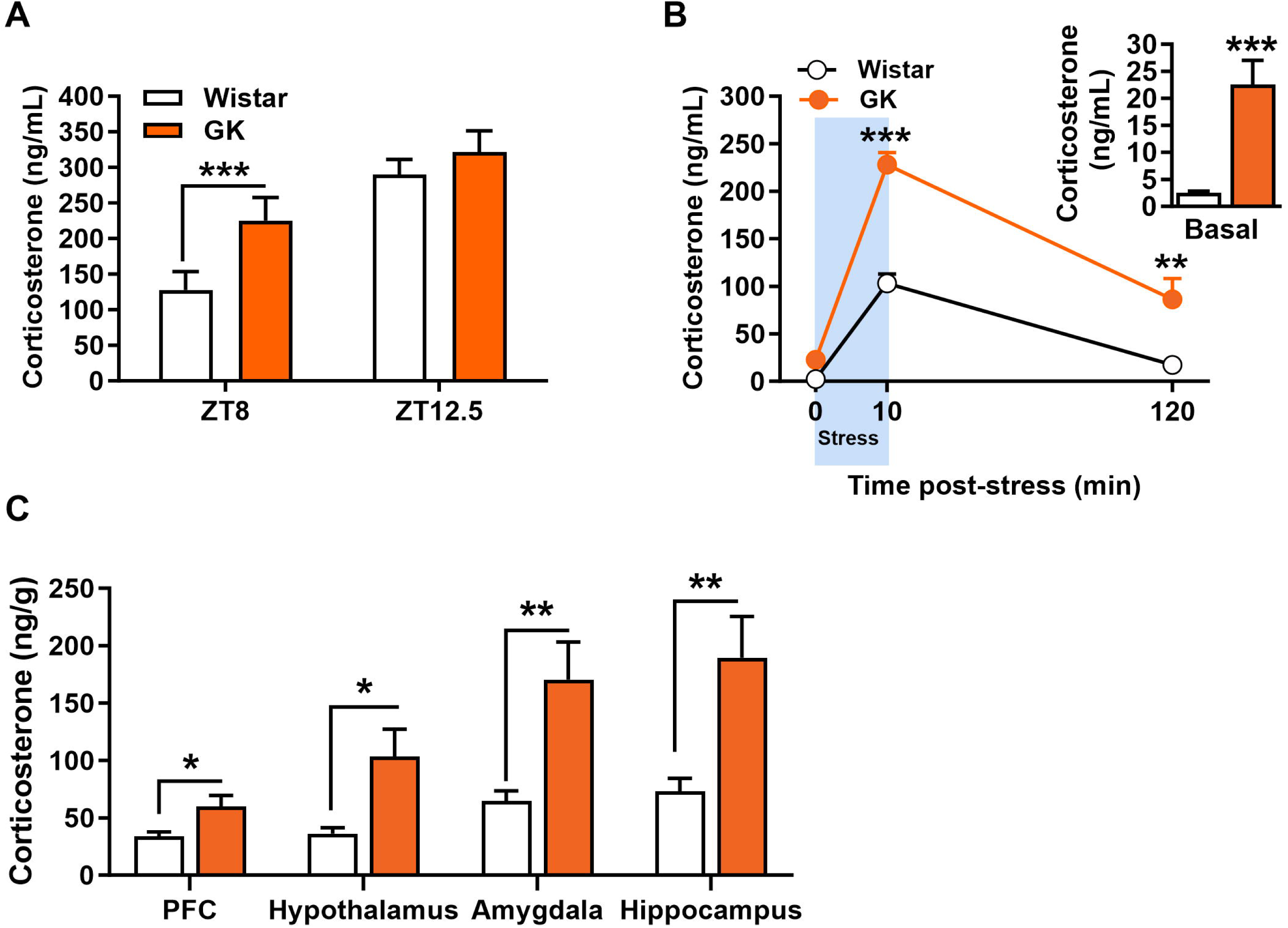
Goto-kakizaki (GK) rats show hypothalamus-pituitary-adrenal axis hyperactivity and enhanced levels of corticosterone in emotion-related brain areas. **(A)** In GK rats, corticosterone levels in plasma were increased in the light phase (zeitgeber time, ZT8) and unaffected at the beginning of the dark phase (ZT12.5) (Wistar, n=9; GK n=5). **(B)** GK rats showed higher basal corticosterone in plasma and enhanced stress-induced corticosterone release (Wistar, n=8; GK n=8). **(C)** GK rats had higher levels of corticosterone in the prefrontal cortex (PFC) (Wistar, n=11; GK n=9), hypothalamus, amygdala, and hippocampus (Wistar, n=9; GK n=10). Data are expressed as mean ± SEM. Two-way ANOVA with repeated measures followed by Tukeýs post-hoc test: (A, B); Unpaired Student’s t-test: (insert B, basal corticosterone) and (C). *p<0.05, **p<0.01 and ***p<0.001 vs Wistar rats.

We then examined whether GK rats exhibit changes in neuronal activation after a mild stress (10 min novelty exposure) using immunostaining of the immediate early gene (IEG) early growth response 1 (EGR1), which has been proposed as an important marker in the context of neuropsychiatric disorders [29]. Brain mapping (**Figure 3**) revealed that GK rats tended to have lower EGR1^+^ cells in the prelimbic (PrL) and infralimbic (IL) areas of the mPFC than Wistar rats (t_(13)_=1.88, p=0.082 and t_(13)_=2.10, p=0.056 respectively; **Figure 3A-D**) while no difference was observed in the cingulate (Cg) area. In addition, GK rats showed higher EGR1^+^cells in response to stress in the (PVN) (t_(13)_=2.34, p=0.036; **Figure 3E-H**), central amygdala (Ce) (t_(11)_=2.36, p=0.038) and basolateral nucleus (BL) of the amygdala (t_(11)_=2.81, p=0.017) (**Figure 3I-N**). GK rats also displayed a tendency for higher EGR1^+^ cells than controls in the dentate gyrus (DG) of the hippocampus (t_(13)_=1.84, p=0.088) while there was no difference between groups in CA1 and CA3 (**Figure 3O-R**). Overall, our data suggest that T2D modifies brain response to a mild stress.

**Figure 3.**
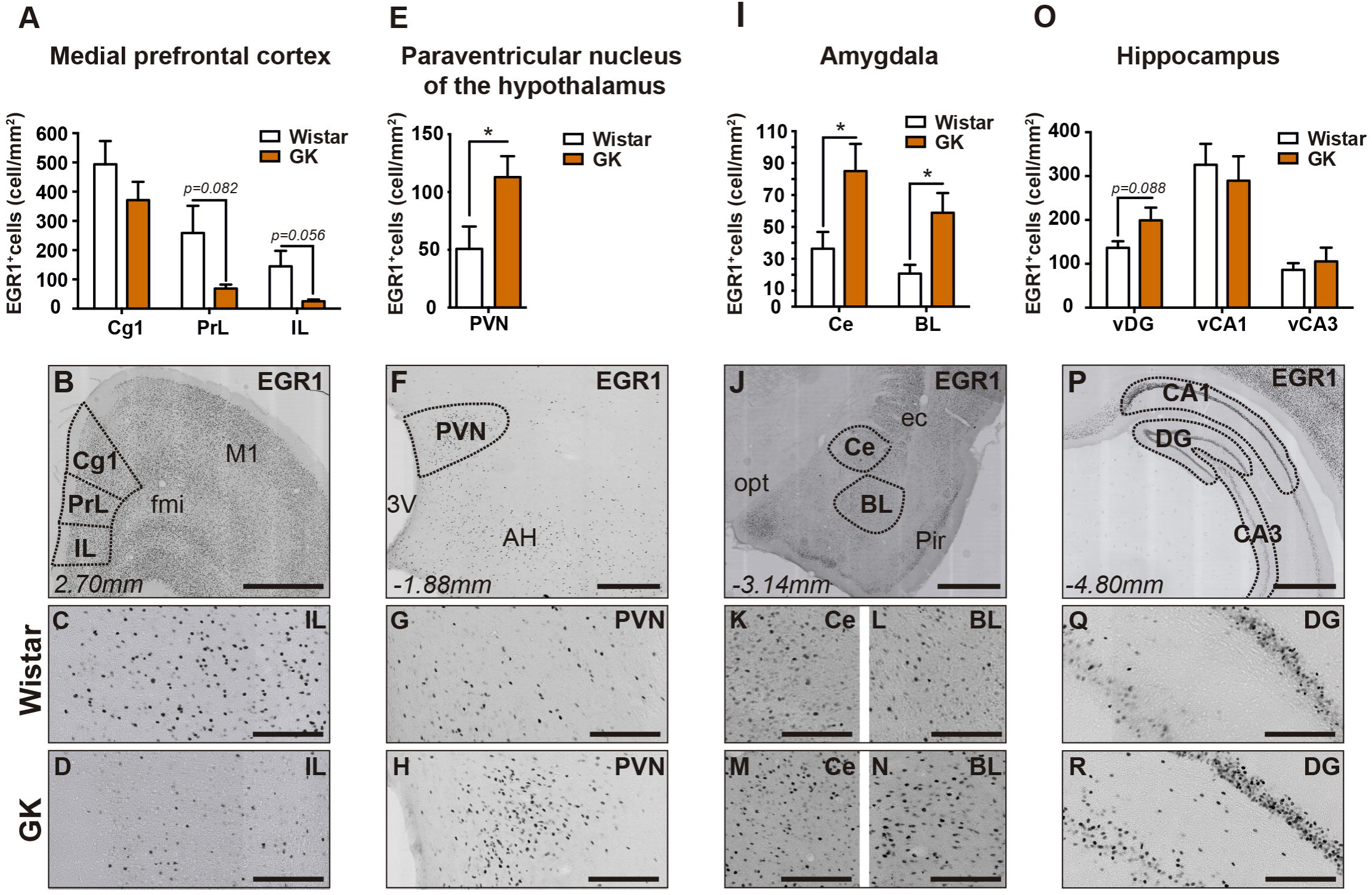
Differential neuronal activity in response to mild stress in Goto-kakizaki (GK) rats. **(A)** GK rats tended to have lower density of EGR1^+^ neurons in prelimbic (PrL) and infralimbic (IL) prefrontal cortex. **(B)** Representative image for EGR1 staining in medial prefrontal cortex. **(C)** Higher magnification of a representative image of the IL prefrontal cortex of Wistar rats and **(D)** GK rats. **(E)** GK rats have higher EGR1^+^ neurons density in the paraventricular nucleus (PVN) of the hypothalamus. **(F)** Representative image for EGR1 staining in PVN. **(G)** Higher magnification of a representative image of the PVN of Wistar rats and **(H)** GK rats. **(I)** Increased EGR1+ neurons density in both, the central and the basolateral amygdala (Ce and BL, respectively) in GK rats. **(J)** Representative image for EGR1 staining in the amygdala. **(K** and **L)** Higher magnification of a representative image of Ce and BL of Wistar rats. **(M** and **N)** Higher magnification of a representative image of Ce and BL of GK rats. **(O)** GK rats tended to have increased density of EGR1^+^ neurons in the dentate gyrus (DG) of the hippocampus. **(P)** Representative image for EGR1 staining in the hippocampus. **(Q)** Higher magnification of a representative image of the DG of Wistar rats and **(R)** GK rats. Scale bars of (B), (F), (J) and (P) correspond to 2mm. Scale bars of (C-D), (G-H), (K-N) and (Q-R) correspond to 500μm. Data are expressed as mean ± SEM, Wistar, n=7; GK, n=8 (Ce, BL, GK n=6; vCA3, Wistar n=6). For each brain area, a blind quantification of EGR1+ cells was conducted in 3-5 sections per animal. Unpaired Student’s t-test: (A), (E), (I) and (O). *p<0.05 vs Wistar rats.

### 3.3 Type 2 diabetes without obesity affects the expression of glucocorticoids signaling, inflammation and serotonin-related genes in the medial prefrontal cortex

Since altered HPA axis and diabetes have been linked to higher neuropsychiatric vulnerability, we hypothesized that GK rats exhibit altered genes expression in anxiety- and mood-related brain areas such as mPFC, ventral hippocampus and amygdala. In the mPFC, we found that GK rats showed an upregulated gene expression of *Nr3c2*, coding for mineralocorticoid receptor (MR) (t_(14)_=2.18, p<0.05) and target genes of the glucocorticoid receptor, namely *Sgk1* (t_(14)_=2.83, p<0.054) and *Ddit4* (coding for REDD1) (t_(14)_=2.04, p=0.061) (**Figure 4A**). Moreover, in the mPFC of GK rats we found a 4-fold increase of *Il-6* transcript levels (t_(13)_=2.99, p<0.05; **Figure 4A**), and reduced *Socs3* gene expression (t_(14)_=2.22, p<0.05; **Figure 4A**), while other inflammation-related genes such as *Il-1*β*, Tnf-*α*, Gp130* and *Stat3* remained unaffected in GK rats (data not shown). In this brain area, the mRNA levels of the serotonin receptor *Htr1a* (t_(14)_=4.35, p<0.001; **Figure 4A**) and the serotonin transporter *Slc6a4* (5HTT) (t_(14)_=2.48, p<0.05; **Figure 4A**) were higher in GK compared with Wistar rats. T2D without obesity only slightly affected genes expression in the other brain areas studied, i.e, the amygdala (**Figure 4B**) and ventral hippocampus (**Figure 4C**). Thus, we found that GK rats exhibited higher mRNA levels of *Nr3c2* (t_(13)_=2.49, p<0.05;) and the serotonin receptor *Htr1a* (5HTR1A) (t_(12)_=2.28, p<0.05) in the amygdala (**Figure 4B**), and higher mRNA levels of the serotonin receptor *Htr2a* (5HTR2A) in the ventral hippocampus (t_(12)_=4.16, p<0.01; **Figure 4C**). In contrast, the expression of inflammation-related genes remained unaffected by the T2D in these brain areas (**Figure 4B** and **4C**).

**Figure 4.**
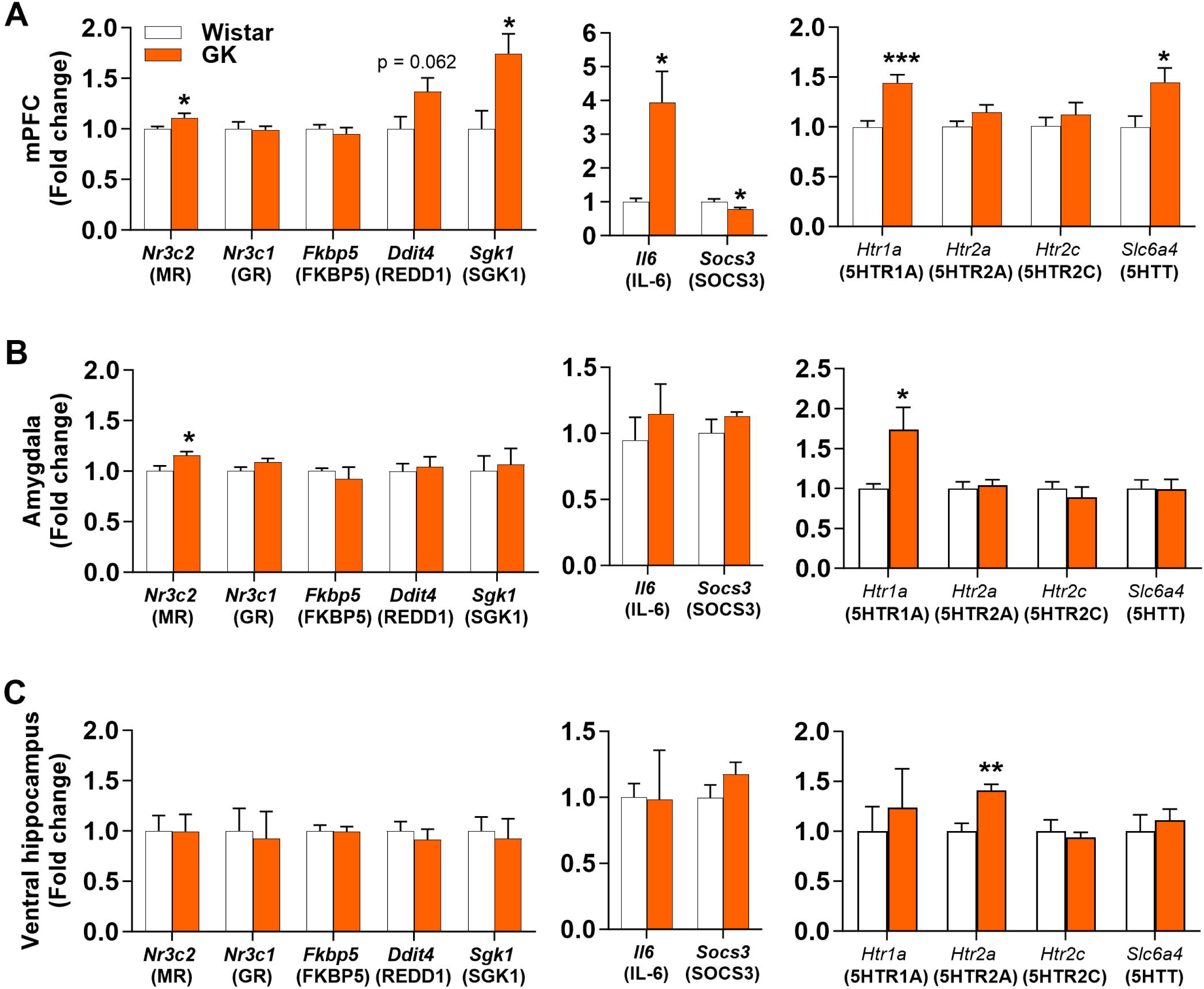
Goto-kakizaki (GK) rats have increased expression of glucocorticoids, inflammation and serotonergic-related genes in the medial prefrontal cortex. Gene expression of glucocorticoids (nuclear receptor subfamily 3 group C member 1 and 2: *Nr3c1, Nr3c2*; FKBP prolyl isomerase 5: *Fkbp5;* DNA damage inducible transcript 4: *Ddit4*; and serum/glucocorticoid regulated kinase 1: *Sgk1*), inflammation (interleukin 6: *Il6* and suppressor of cytokine signaling: *Socs3*) and serotonergic (5-hydroxytryptamine receptor 1A, 2A and 2C: *Htr1a*, *Htr2a* and *Htr2c*; solute carrier family 6 member 4: *Slc6a4*)-related genes in **(A)** medial prefrontal cortex (mPFC), **(C)** amygdala, and **(D)** ventral hippocampus. Data are expressed as mean ± SEM, n= 5-8 rats *per* group (details in supplemental table S2). Unpaired Student’s t-test. *p<0.05, **p<0.01 and ***p< 0.001 vs Wistar rats.

### 3.4 Type 2 diabetes without obesity alters neuronal morphology and reduces the astrocyte density in the medial prefrontal cortex

Our data highlight that T2D without obesity targets the mPFC, by enhancing glucocorticoid, inflammation and serotonergic-related pathways. Since the architecture of this brain area is particularly affected by overexposure to stress or to high glucocorticoid levels, we examined pyramidal neurons morphology and astrocytes density in GK rats [30]. GK rats showed a reduced complexity of the apical and basal dendritic arborizations in comparison with Wistar controls (**Figure 5A**). Sholl analysis revealed that apical dendritic length significantly varied with the distance from soma (two-way ANOVA with repeated measures, F_(20, 860)_=33.08, p<0.0001) which tended to be altered by genotype (F_(1,43)_=3.21, p=0.08) (**Figure 5A**). The reduced dendritic length of GK rats was particularly marked in the proximal part of the apical tree (0-260 µm from soma, t_(43)_=2.33, p<0.05; **Figure 5B**). In the basal arbor, an interaction between the genotype and the distance from the soma was identified on dendritic length (F_(11,440)_=4.13, p< 0.0001, **Figure 5A**). Planned comparisons indicated that GK rats had reduced dendritic length at 60, 80, 100 and 120 µm from the soma (p at least<0.05; **Figure 5A**). The total dendritic length of the basal arborization was also markedly reduced in GK rats (t_(43)_=4.59, p<0.001; **Figure 5B**). The spine density in the basal arborization of layer II/III pyramidal neurons in the mPFC was reduced in GK rats (t_(208)_ = 1.94, p<0.05; **Figure 5C**), whereas their apical tree remained unaltered (**Figure 5C**). Chronic stress and psychiatric conditions alter astrocytes [31–33]. Here we showed that the number of astrocytes were reduced in IL of GK rats (t_(7)_=4.52, p<0.01; **Figure 5D**) with a similar tendency in PrL (t_(7)_=2.27, p=0.057; **Figure 5D**).

**Figure 5.**
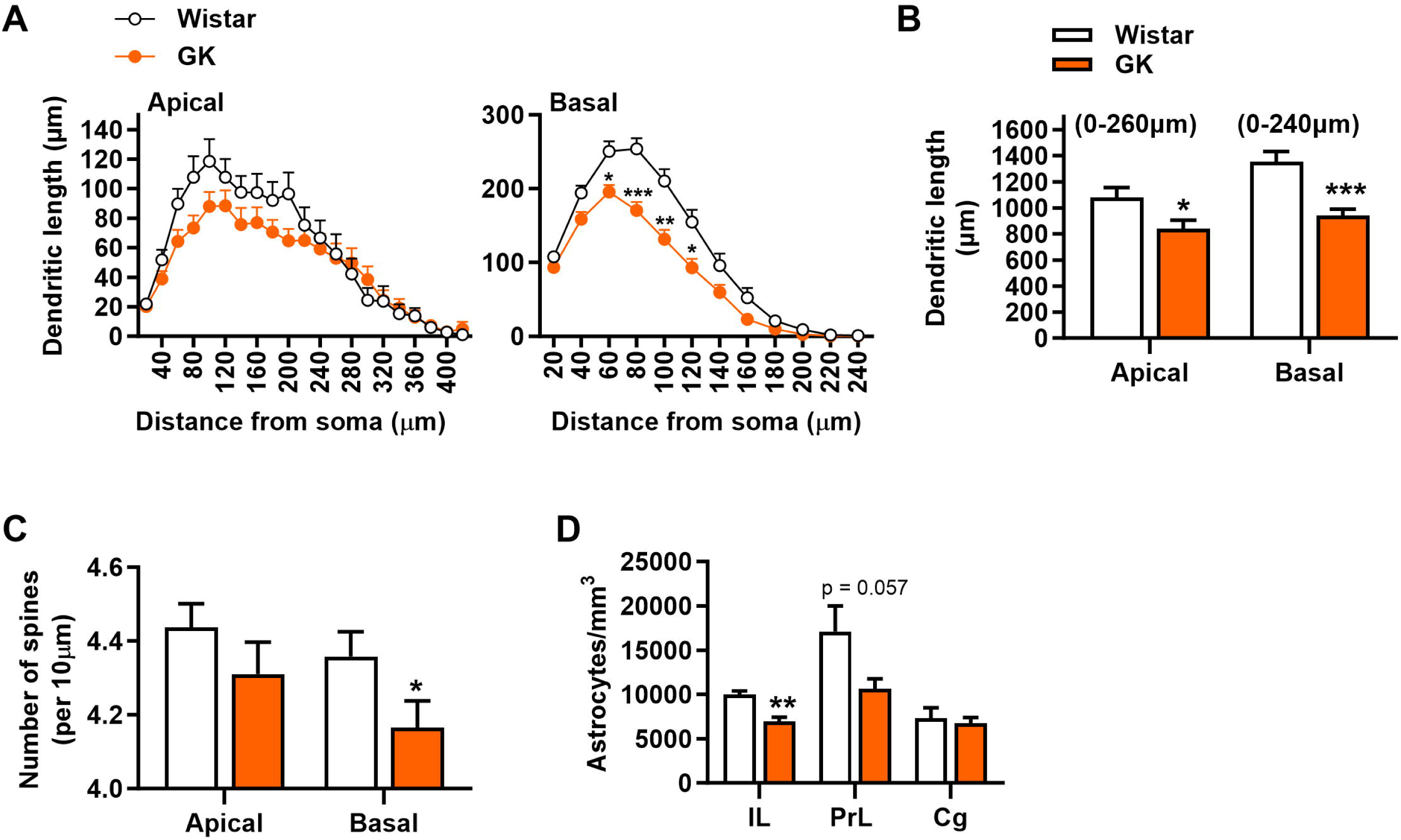
The Goto-kakizaki (GK) phenotype alters the morphology of neurons and reduced astrocyte density in the medial prefrontal cortex. **(A)** GK rats had diminished length of basal dendrites of pyramidal neurons located at different distances from the soma in the medial prefrontal cortex (mPFC) layer II/III. **(B)** Both, apical and basal total dendritic length of pyramidal neurons from the mPFC layer II/III were reduced in GK. **(C)** The pyramidal neurons of GK rats in mPFC layer II/III showed reduced number of basal dendritic spines. **(D)** GK rats showed lower astrocyte density in the infralimbic (IL) PFC and a similar tendency in the prelimbic (PrL) PFC. Data are expressed as mean ± SEM. The blind analyses were conducted in 4-6 neurons per animal in each mPFC subregion, in the case of dendritic morphology and spine density (Wistar, n=4; GK n=5; n=20-26 neurons per groups for dendritic morphology and 99-127 segments per groups for spine density); and 3-5 sections per animal (Wistar, n=4; GK n=5) in each mPFC subregion, in the case of astrocytes density. Two-way ANOVA with repeated measures followed by planed comparisons: (A); Unpaired Student’s t-test: (B-D). *p<0.05, **p<0.01 and ***p<0.001 vs Wistar rats.

### 3.5 Type 2 diabetes without obesity induces exacerbated anxiety, reduced social interaction and increased depressive-like behavior

Given the key role of diabetes, stress and PFC abnormalities in anxiety and mood disorders, we hypothesized that GK rats will exhibit emotional alterations. We studied GK rats’ emotional behaviors in different validated paradigms for anxiety-like and depressive-like behavior such as light/dark test, open-field, tunnel test and FST. We also examined their social behavior and their circadian motor activity which are known to be affected in humans suffering from anxiety disorders and major depression.

In the light-dark test, GK rats spent less time (t_(14)_=4.30, p<0.001; **Figure 6A**) and did less visits (t_(14)_=3.47, p<0.01; **Figure 6B**) to the light compartment than Wistar controls. Moreover, the time spent in the center area of the open-field (t_(12)_=2.72, p<0.05; **Figure 6C**) and the number of visits to this area (t_(12)_=3.56, p<0.01; **Figure 6D**) were reduced in GK rats. GK rats also exhibited an increased anxiety-like behavior in a familiar environment since spent more time in the familiar tunnel placed as enrichment in their home cage (t_(15)_=8.51, p<0.001; **Figure 6F**) and did higher number of visits to the tunnel than Wistar rats (t_(15)_=5.69, p<0.001; **Figure 6F**), although the latency to enter in the tunnel was not affected by the genotype (Wistar:

**Figure 6.**
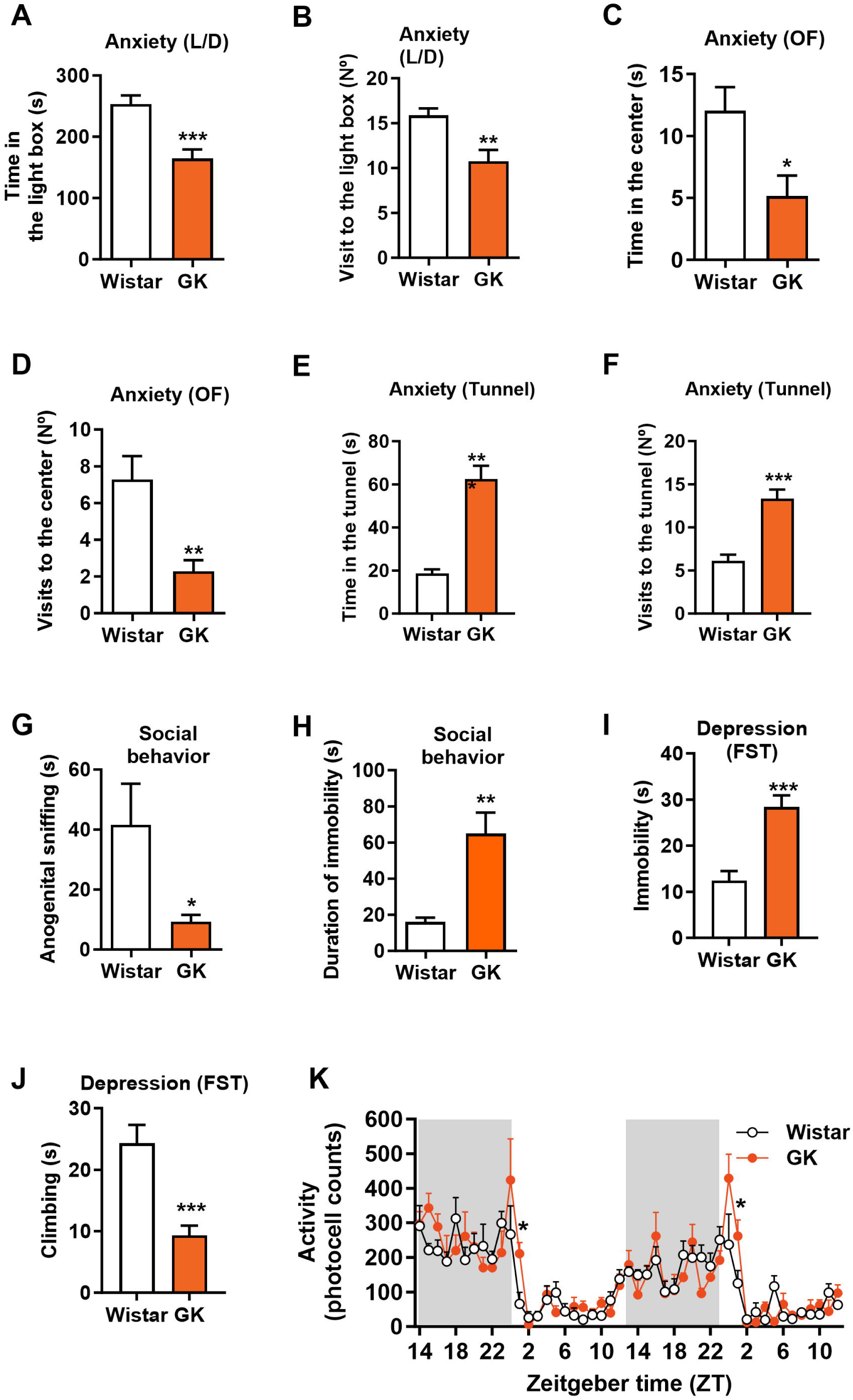
Goto-kakizaki (GK) rats exhibit an exacerbated anxiety and depressive-like behavior and an altered social behavior and circadian locomotor activity. **(A)** In the light-dark test (L/D), GK rats spent less time in the light box and **(B)** did less visits to the light compartment than Wistar rats (Wistar, n=8; GK n=8). **(C)** The time spent in the center of the open-field (OF) was lower in GK rats **(D)** that also did less visits to this area (Wistar, n=7; GK n=7). **(E)** GK rats had increased time spent in the tunnel **(F)** and did more visits to the tunnel **(G)** (Wistar, n=11; GK n=6). In the social interaction test, GK rats did less anogenital sniffing and **(H)** showed a higher time spent immobile (Wistar, n=11; GK n=6). In the forced-swimming test (FST), **(I)** GK rats are more immobile during the first minute. **(J)** Climbing behavior during the first minute of the FST was also reduced in GK rats (Wistar, n=13; GK n=13). **(K)** Circadian locomotor activity. At the beginning of the light phase GK rats showed higher activity (Wistar, n=7; GK n=6). Zeitgeber time (ZT) 0 indicates lights ON. Data are expressed as mean ±SEM. Unpaired Student’s t-test: (A-J); Two-way ANOVA with repeated measurements followed by planned comparisons: (K). *p<0.05, **p<0.01 and ***p<0.001 vs Wistar rats.

7.12 (s) ±4.9 and GK: 8.00 (s) ±4.9). GK rats showed reduced social investigative behavior, as indicated by the decrease of the anogenital sniffing duration (t_(14)_=2.30, p=0.0371; **Figure 6G**), combined with an increase of their immobility during the social interaction test (t_(14)_=4.10, p=0.0011; **Figure 6H**). During the pretest of the FST, GK and Wistar rats showed the same level of immobility demonstrating an intact motor function in GK rats (t_(24)_=1.53, p=0.14, data not shown). During the test session, two-way ANOVA revealed a significant genotype x time interaction on immobility (F_(4,120)_=10.47, p<0.0001) and climbing (F_(4,120)_=7.52, p<0.001). During the first 3 min of the test, GK rats were more immobile than Wistar rats (t_(24)_=4.92, p<0.001, **Figure 6I**) and had reduced climbing behavior (t_(24)_=4.42, p<0.001, **Figure 6J**), indicating higher passive coping and lower active coping, respectively. As expected, locomotor activity was higher during the dark phase in comparison with the light phase (light-dark effect, F_(1,11)_= 259, p<0.0001; **Figure 6K**). However, while no change in overall locomotion was noted between groups, GK and Wistar rats showed a different changes in their locomotion across the circadian rhythm (genotype x time interaction, F_(46,506)_=1.78, p=0.001; **Figure 6K**). Planned comparisons revealed that GK rats had higher locomotor activity at the beginning of the inactive light phase (day 1, ZT1, F_(1,11)_=9.53, p<0.01; day 2, ZT1, F_(1,11)_=5.54, p=0.038), suggesting a shifted of their circadian rhythm. These results demonstrated that T2D without obesity elicits social interaction deficits, and exacerbated anxiety, in both novel and familiar contexts, and depressive-like behaviors coupled with altered circadian rhythms.

### 3.6 The normalization of plasma glucocorticoids levels in Goto-kakizaki rats partially improves metabolic and emotional abnormalities associated with diabetes

In order to identify potential mechanisms involved in the impaired emotional behavior in GK rats, we examined the role of glucocorticoids. Thus, we normalized peripheral glucocorticoids levels in GK rats by conducting an adrenalectomy in combination with low-dose corticosterone replacement in the drinking water to reproduce circadian physiological plasma corticosterone levels.

Contrary to GK sham rats, adrenalectomized GK (GK ADX) rats showed a marked reduction of plasma corticosterone levels and no corticosterone rise after stress (two-way ANOVA, group x stress, F_(1,14)_=34.07, p<0.0001; Tukeýs post-hoc test, GK ADX rats, basal versus stress, n.s ; GK sham rats, basal versus stress, p<0.001, **Figure 7A**). Moreover, peripheral corticosterone normalization in GK rats reduced corticosterone levels in the PFC compared with sham-operated GK rats (t_(14)_=4.04, p<0.01; **Figure 7B**). Glycemia differently varied across GK groups in response to the stress. Indeed, an interaction between the GK groups x stress was identified (F_(1,14)_=7.50, p<0.05 **Figure 7C**). Glycemia was increased in response to a mild stress in sham-operated GK rats (p<0.001 versus basal) but not in adrenalectomized GK rats (n.s. versus basal). In addition, GK ADX rats showed lower glycemia after stress than GK sham (p<0.001). The analysis restricted to basal condition also revealed a tendency in GK ADX rats for an improvement of their hyperglycemia (t_(14)_=1.88, p=0.08). On the other hand, adrenalectomized GK rats have higher insulin levels in plasma than sham-operated GK rats (t_(14)_=3.31, p<0.01 **Figure 7D**).

**Figure 7.**
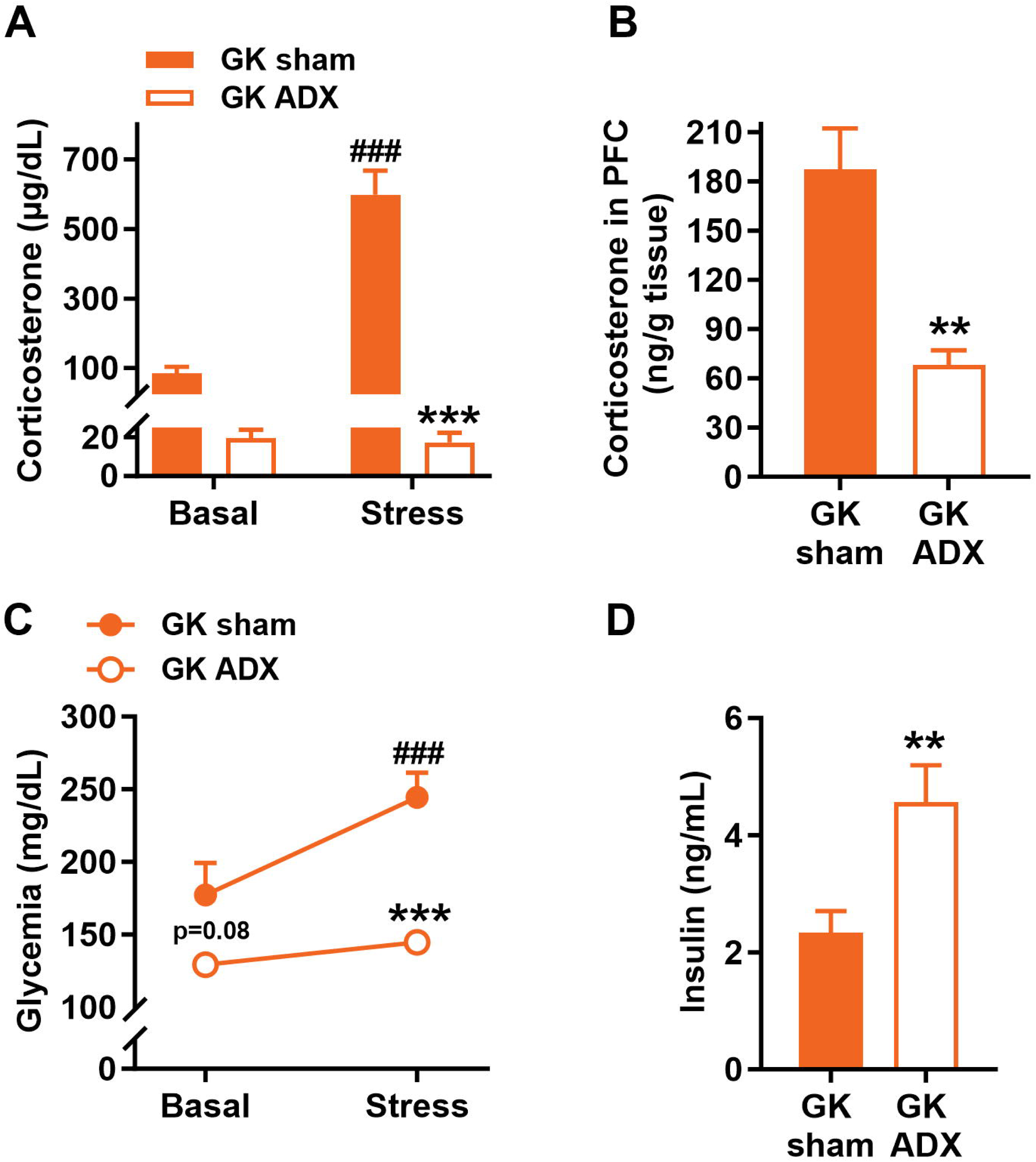
Normalization of glucocorticoids improves glycemia in Goto-kakizaki (GK) rats. **(A)** Compared with sham-operated GK rats, adrenalectomized GK rats (GK ADX) had reduced corticosterone in both, basal conditions and in response to stress and **(B)** lower corticosterone levels in prefrontal cortex. **(C)** GK ADX had reduced glycemia in basal conditions and after stress and glucocorticoids normalization prevented the increased glucose levels in plasma after stress. **(D)** Insulin levels in plasma were higher in GK ADX than in GK sham. Data are expressed as mean ±SEM, Wistar, n=9; GK n=7 (corticosterone in PFC, GK n=6). Two-way ANOVA with repeated measures followed by Tukeýs post-hoc test: (A) and (C); or Unpaired Student’s t-test (B and D). *p<0.05, **p<0.01 and ***p<0.001 vs GK sham and ^###^p<0.001 vs basal levels.

In the light-dark test, adrenalectomized GK rats showed a tendency to do more visits to the light box (**Figure 8A**, two-way ANOVA with repeated measures; Group effect: F_(1,14)_=3.60, p=0.07) than sham-operated GK rats, while the time spent in this compartment remained unaffected (**Figure 8B**). During the test, the exploration of the light compartment was very low during the first 5min period, but increased during the second 5min period. An interaction between GK groups x period was identified on the number of visits to the light box (F_(1,14)_=5.78, p<0.05; **Figure 8A**). Adrenalectomized GK rats increased their visits in the second period (Tukeýs post-hoc, p<0.01 period 2 versus period 1), doing more visits than sham-operated rats in this period (Tukeýs post-hoc, p<0.05 GK ADX period 2 versus GK sham period 2, **Figure 8A**). In the open-field, adrenalectomized GK rats showed a tendency to spent more time in the center than sham-operated GK rats (t_(14)_=1.95, p=0.07; **Figure 8C**), and travelled a higher distance in the center (t_(14)_=2.14, p<0.05; **Figure 8D**). Adrenalectomized GK rats had less inactivity in the periphery of the open-field (t_(14)_=3.39, p<0.01; **Figure 8E**). Adrenalectomized GK rats also exhibited an improvement of their anxiety in a familiar context since their latency to enter in the tunnel was increased in comparison with sham-GK (t_(14)_=2.17, p<0.05.; **Figure 8F**) although the number of visits to the tunnel remained unchanged (**Figure 8G**). Adrenalectomized GK rats spent more time in contact with the juvenile rat compared with sham-operated rats (t_(13)_=2.88, p<0.05; **Figure 8H**) showing that the normalization of glucocorticoids improves the social behavior of GK rats. Both groups showed similar immobility during the first period of the FST, suggesting that the restoration of glucocorticoids did not attenuate depressive like behavior (**Figure 8I**).

**Figure 8.**
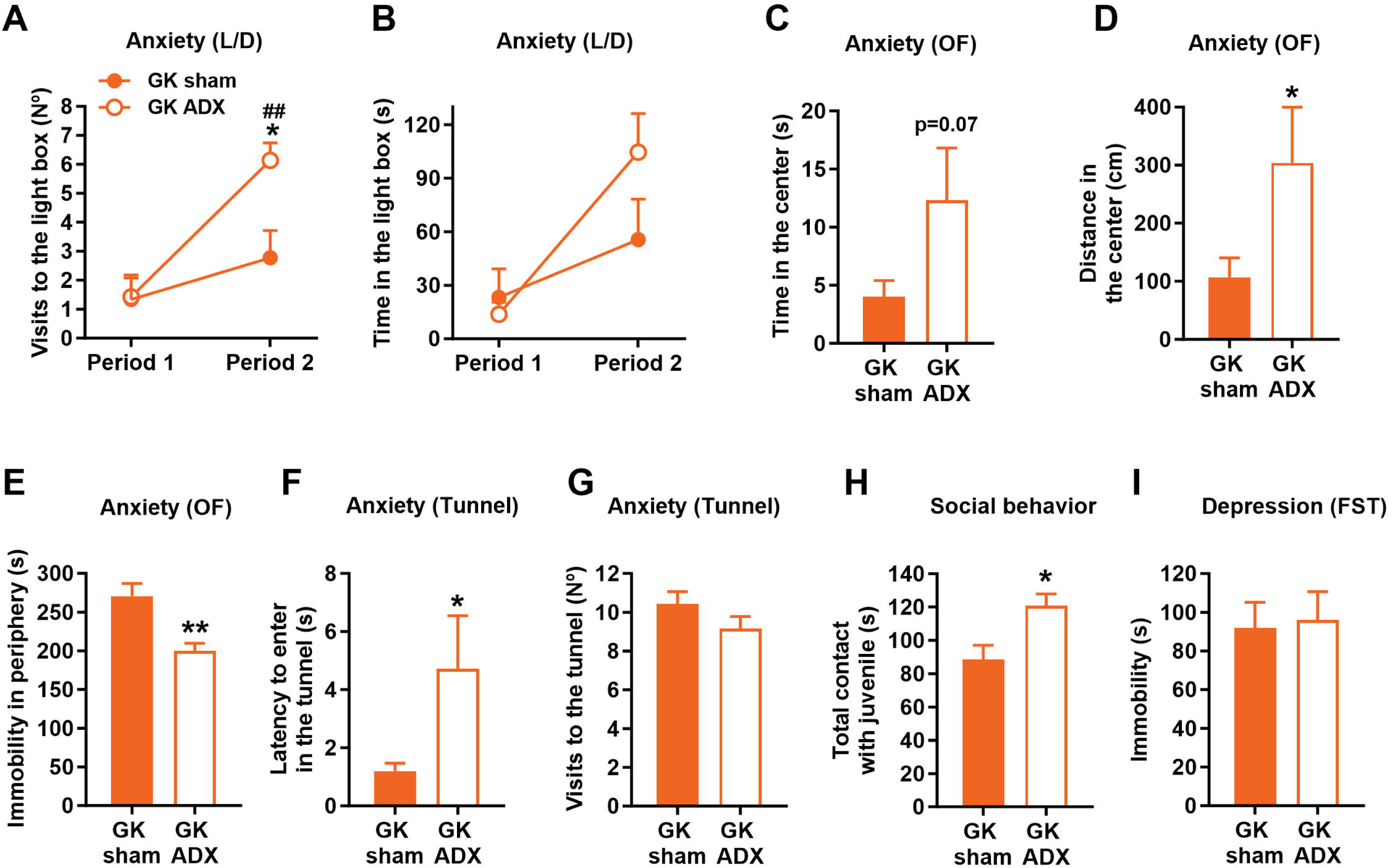
Normalization of glucocorticoids attenuates anxiety but not depressive-like behavior in Goto-kakizaki (GK) rats. **(A)** In the light-dark test (L/D), GK rats with normalized glucocorticoids levels (GK ADX) did more visits to the light box during the second period of the light-dark test than sham-operated rats while **(B)** the time spent in this compartment remained unaffected. **(C)** In the open-field test (OF), GK ADX tended to spend more time in the center, **(D)** significantly travelled more distance in the center, and **(E)** were less immobile in the periphery of the open-field. **(F)** GK ADX had increased latency to enter in the tunnel but, **(G)** the number of visits to the tunnel remained unchanged compared with sham-operated rats. **(H)** GK ADX spent more time in contact with juvenile in the social interaction test. **(I)** In the forced swim test (FST), immobility time of GK ADX rats was similar to that of GK sham. Data are expressed as mean ±SEM, Wistar, n=9; GK n=7 (social interaction, Wistar, n=8). Two-way ANOVA with repeated measures: (A-B); or Unpaired Student’s t-test (C-I). *p<0.05 and **p<0.01 vs GK sham and ^###^p<0.001 vs period 1.

Overall, our data suggest that inhibiting the corticosterone hypersecretion in non-obese T2D rats may improve anxiety and social behaviors but not the depressive-like behavior.

## 4. Discussion

Obesity is a well-established risk factor for brain and mental health issues, particularly depression [34]. The global rise in type 2 diabetes (T2D) incidence has largely been attributed to the obesity epidemic. However, the prevalence of T2D in individuals with a lean or normal body mass index has recently surged, particularly in certain populations such as those in Asia [35,36]. While obesity and T2D are both linked to profound physiological changes, these alterations are not necessarily identical between the two conditions [37]. Epidemiological studies investigating the relationship between T2D and depression highlight a complex interplay between diabetes, obesity, and depression risk. Some studies suggest that obesity and elevated fasting blood glucose levels may actually be associated with a reduced risk of depression [38,39]. In contrast, a prospective study reveals that a prediabetic state, rather than high central adiposity, predicts the onset of major depression [40]. In this context it remains unclear it remains unclear whether T2D independently increases the risk of neuropsychiatric symptoms in lean individuals.

Preclinical studies often use rodent models exposed to chronic high-fat diets as a clinically relevant method to study T2D. However, these models provide limited insights into the effects of T2D independent of obesity. The Goto-Kakizaki (GK) rat, a non-obese T2D model, offers a valuable alternative for studying the pathophysiology of non-obese T2D. This model exhibits key T2D characteristics in both rats and humans, such as pancreatic β-cell dysfunction and impaired insulin secretion. The GK model has enhanced our understanding of the less explored factors contributing to the etiology of T2DM.These mechanisms include complex interactions between polygenic inheritance and early-life epigenetic programming due to exposure to a hyperglycemic maternal environment [41]. The GK rat model allows to disentangle emotional behavior changes directly linked to T2D from those caused by obesity-related metabolic complications or hypercaloric diets. In this study, we demonstrated that T2D in lean GK rats leads to a depressive phenotype and heightened anxiety. Our results are consistent with previous works showing that T2D in mice and rats overweighed models are associated with a marked increase of anxiety-like behavior in different paradigms such as elevated-plus maze, open-field and novelty suppressed feeding test [9,12–14,16,42,43]. Additionally, we demonstrated that hyper-anxiety in GK rats is seen not only in typical stress paradigms such as light-dark test, open-field but is also detected in non-stressful home cage environment such as in the tunnel test. Regarding depressive-like behaviors, previous studies in overweight T2D models have yielded mixed results. Some report no significant effects in the tail suspension test (TST) or forced swim test (FST) [12,13,42] while others find impairments in tests like the splash test, sucrose preference test, and female urine test [16,42]. Additional studies, found however an increase of depressive-like behavior in the FST in diabetic db/db mice [44] and in mice exposed to 12 week regimen of high-fat diet [10]. In GK rats, we observed pronounced depressive phenotypes, including high immobility in the FST, reduced social behavior, and disrupted circadian locomotor rhythms, consistent with depression models [45,46]. Our data confirm and extend previous studies showing behavioral alterations in GK rats [21,47] highlighting the GK model as a robust tool for investigating anxiety and depression associated with diabetes. Our findings also expand on previous results from the streptozotocin-induced T1D model, which similarly exhibits heightened anxiety and depressive-like behaviors [48]. However, our data contrast with a prior study showing that 16 weeks of fructose exposure—a nutritional model of lean T2D—had no effect on emotionality [42]. Exposure to diabetes during critical developmental periods may also contribute to the pronounced effects of T2D on the brain and behavior in GK rats. Specifically, GK embryos develop in a diabetic prenatal environment, and GK offspring are raised by diabetic mothers. Additionally, impaired maternal care in GK dams may further exacerbate stress-related brain and behavioral phenotypes. Overall, these findings demonstrate that obesity is not a necessary factor for the significant intensification of anxiety- and depressive-like behaviors.

Despite the extensive characterization of T2D pathology in GK rats, neurobiological alterations have not received as much attention. Our findings reveal that GK rats not only exhibit basal hyperglycemia, basal hyperinsulinemia, and impaired glucose-induced insulin secretion but also show elevated corticosterone levels in both the brain and plasma during the resting phase. Additionally, they display an exaggerated HPA-axis response to mild stress. Previous studies have also reported associations between HPA-axis hyperfunction and T2D 37. Although it remains unclear whether HPA-axis dysregulation is a cause or consequence of T2D in GK rats, it likely exacerbates the diabetic phenotype and its associated complications. In our study, HPA-axis hyperactivity appeared to contribute to certain emotional disturbances in GK rats. Adrenalectomized GK rats showed significantly reduced corticosterone levels in the prefrontal cortex (PFC) and partial improvements in anxiety and social behaviors. However, depressive-like behaviors in the FST persisted. Since adrenalectomy also improved blood glucose and insulin levels in GK rats, the effects on emotional behaviors may be mediated, at least in part, by metabolic improvements. These findings align with prior research indicating that normalizing glucocorticoid levels can alleviate diabetes-related cognitive deficits [26].

Remarkably, GK rats exhibit brain abnormalities reminiscent of those induced by early-life stress, chronic stress, or prolonged exposure to high levels of glucocorticoids [28,30,49]. Using immediate early gene (IEG) immunostaining, we observed increased neuronal activation in the amygdala and paraventricular nucleus (PVN) of GK rats, along with reduced activation in the infralimbic (IL) region of the medial prefrontal cortex (mPFC) in response to mild stress. These findings suggest impaired communication between the neural circuits involved in emotional processing and stress responses. Notably, amygdala hyperactivity and mPFC hypofunction have been observed in individuals with anxiety disorders and depression, as well as following early-life adversity. [50,51]. The mPFC plays a central role in stress regulation, receiving inputs from the amygdala, hippocampus, and hypothalamus, and exerting inhibitory control over the amygdala and PVN to regulate the HPA axis [52]. Indeed, the ventral mPFC exerts inhibitory control over amygdala activity in response to aversive stimuli in humans [52]. Glutamatergic neurons in the mPFC have been involved in the inhibitory regulation of the amygdala and PVN through activation of inhibitory neurons in the bed nucleus of the stria terminalis contributing to the negative feedback of the HPA axis [53,54]. Structural changes in mPFC have been hypothesized to contribute to depressive disorders [55]. A large literature both in humans and in animal models demonstrates that depression, chronic stress or chronic glucocorticoids administration are associated with neuronal architecture remodeling and reduced glial cells density in mPFC [30,56,57]. In the present work, we reported reduced neuronal spines density and decreased complexity of the dendritic arborization of pyramidal neurons of the mPFC. We also showed a reduced density of astrocytes in the infralimbic and prelimbic PFC. In line with this, a previous study from Banasr and Duman [31] demonstrates that unpredictable chronic stress in rats reduced the density of astrocytes in the PrL cortex and that glial destruction in PFC led to a depressive-like phenotype.

Our data also revealed an upregulation of the gene expression of glucocorticoid-related transcriptional markers such as REDD1 and SGK1 and the pro-inflammatory cytokine IL-6 in the mPFC which can contribute to reduce the altered synaptic plasticity and to emotional abnormalities reported in GK animals. In line with this finding, SGK1 has been involved in the cortisol-induced decrease in hippocampal neurogenesis and SGK1 mRNA expression is increased in blood of patients with major depression [58]. Similarly, IL-6, in pathological conditions, may affect hippocampal plasticity and is correlated with major depression [59]. Finally, the increase of 5HT1A mRNA expression in the mPFC and amygdala is consistent with previous findings showing higher 5HT1A levels in these brain areas in preclinical model of depression such helpless mice [60] and prenatal stress [61]. Additionally, higher brain 5HT1A receptors in humans has been proposed as a biological trait of major depressive disorder [62]. An important mediator of the effects reported in GK rats may be REDD1, a stress-induced protein acting as an inhibitor of the AKT/mTOR signaling pathway, which has been implicated in various diseases including depression and diabetes [63]. Previous preclinical studies in rodents have shown that elevated glucocorticoid levels and chronic stress are linked to increased REDD1 expression in the mPFC. Overexpression of REDD1 in this brain region leads to synaptic loss and exacerbates anxiety-like and depressive-like behaviors [64]. Notably, REDD1 mRNA expression is also upregulated in the PFC of patients with major depressive disorder [64]. Future research should aim to uncover the potential causal role of REDD1, SGK1, IL-6, and serotonergic signaling in the emotional deficits observed in GK rats.

## 5. Conclusions

Type 2 diabetes (T2D) without obesity induces brain and behavioral phenotypes that closely resemble the effects of chronic stress. Both T2D and depression increase the risk of developing age-related diseases that impair cognition, such as Alzheimer’s disease [65]. These findings underscore the critical need to investigate and develop preventive and therapeutic strategies to address T2D-related complications at early stages. In this context, our study emphasizes the importance of preventing chronic stress in patients with T2D and highlights the potential benefits of targeting glucocorticoid signaling to alleviate the neuropsychiatric symptoms associated with T2D.

## CRediT authorship contribution statement

Conceptualization and experimental design: M. Darnaudéry, B Portha, J. Turner, J Movassat; Investigation: M. Henry, AL. Lépinay, M, Romani-Pérez, K Buyannemekh, M Rincel, A L. Lépinay, D Bailbé, J Brossaud, L Prado, C Pajot; Data analysis: M. Henry, M. Romani-Pérez, M. Darnaudéry; Manuscript writing: M. Henry, M. Romani-Pérez, and M. Darnaudéry. All of the authors have read, revised and approved the final manuscript.

## Availability of data and material

Data will be made available on request.

## Acknowledgements

The authors thank the staff from the animal facility of Nutrineuro lab for animal care. The microscopy was done in the Bordeaux Imaging Center a service unit of the CNRS-INSERM and Bordeaux University, member of the national infrastructure France BioImaging supported by the French National Research Agency (ANR-10-INBS-04). The help of Sebastien Marais is acknowledged.

---

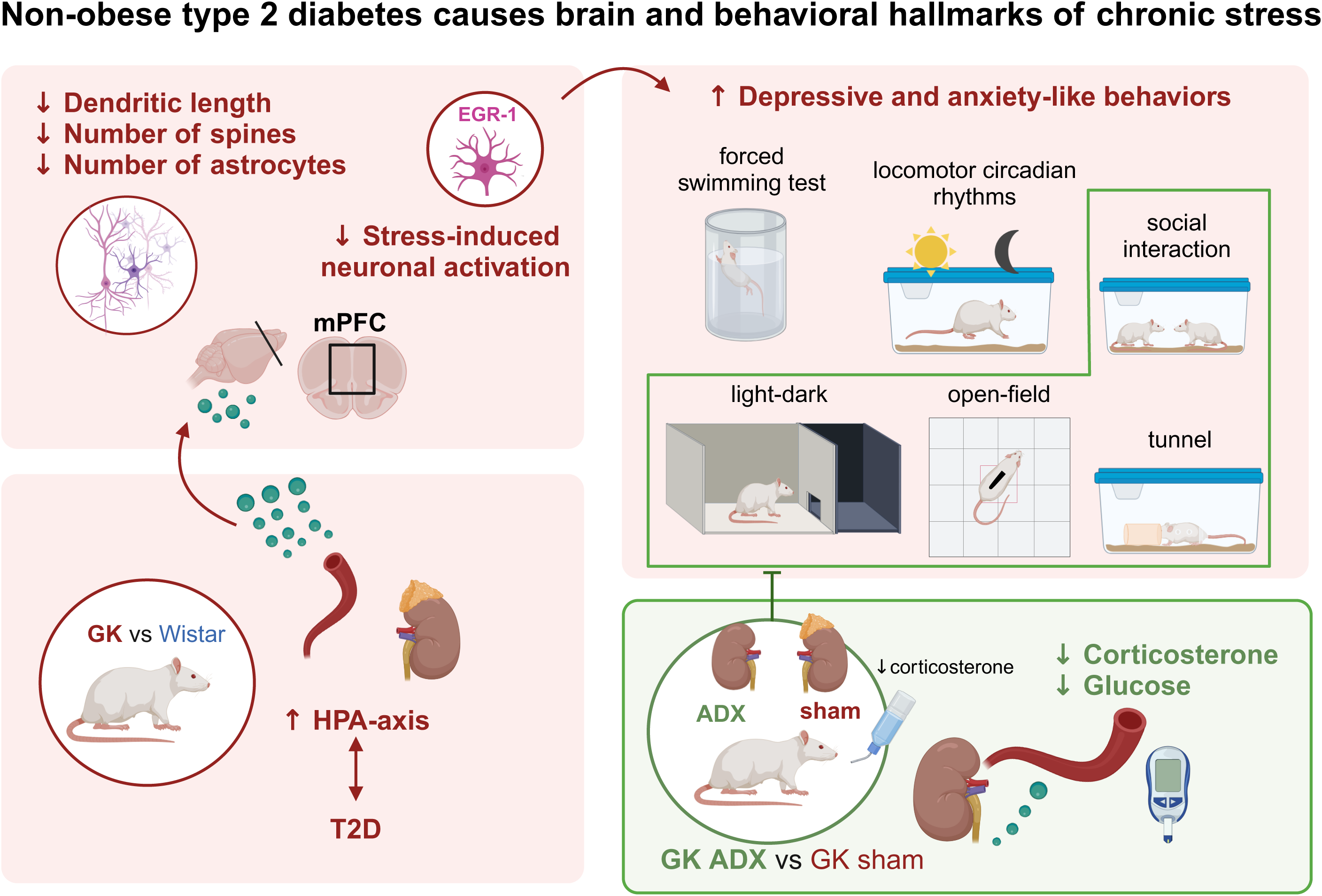

