## Supplementary material for "Non-obese genetic type 2 diabetes causes brain and behavioral hallmarks of chronic stress": SUPLEMENTAL MATERIAL

**Supplementary methods**

**Animals**

Wistar and Goto Kakizaki (GK) female and male rats provided by B2PE, unit BFA, Université Paris Cité and CNRS (Paris, France) were bred in the animal facility at Nutrineuro lab (INRA UMR1286, Bordeaux). After mating, pregnant rats were singly housed during gestation and lactation until weaning. Only male Wistar and GK offspring was used in the present study (a total of 9 cohorts), females were used for other protocols. Animals were group-housed except for the circadian locomotor activity. The collection of biological samples and measurements followed an alternating order between experimental groups. For the measurements, codes were assigned to the animals; the experimental conditions of these codes were revealed during the statistical analysis

**Glycemia, glucose tolerance test and insulin secretion**

Glycemia was measured with ACCU-CHEK performa system (Roche Diagnostics, Mannheim, Germany) in blood collected from the lateral tail vein. The intraperitoneal glucose tolerance test (IPGTT) was conducted by administering glucose (1g/kg) intraperitoneally to 6h-fasted rats followed by blood collection before and 15, 30, 60, 90 and 120 min after glucose administration. Glycemia during the IPGTT was determined in all time points with ACCU-CHEK performa system. Insulin was measured in plasma from blood collected at 0, 15, 30 and 90 min with an ELISA kit (ALPCO, Salem, NH, USA).

**Corticosterone measurements in the brain**

Plasma corticosterone was measured with an ELISA kit (Immunodiagnostic systems, Boldon, United Kingdom) or by in-house radioimmunoassay [1]. In isolated prefrontal cortex (PFC), hypothalamus, amygdala and hippocampus, corticosterone was measured using liquid chromatography–mass spectrometry (LC/MS) (Prominence liquid chromatography system, Shimadzu, Nakagyo, Japan; 5500 Qtrap detector, Sciex,Framingham, USA) as previously described [2]. In brief, animals were deeply anesthetized with halothane and killed by decapitation. PFC, hypothalamus, amygdala and hippocampus were quickly dissected, weighted and stored at -80°C until use. Weighted samples were ultrasonicated with acetonitrile:acetic acid (110:1 v/v) solution and then centrifuged (4500 rpm at 20 °C, 10 min) to collect supernatants. Lipids were removed by incubating samples with hexane (10 min with agitation) and then centrifuged (3500 rpm at 20 °C, 5 min) to collect the lipid-free upper phase. Samples were passed through a Captiva plate (Agilent, Les Ulis, France), evaporated in a heated water bath (40 °C) under nitrogen flow, and re-dissolved in methanol:water solution (50:50 v/v). Deuterated cortisol (500 ng/ml) was added to samples as internal standard (Steraloids Newport, USA). Peaks area ratios of corticosterone were determined through LC/MS (5500 QTRAP, Sciex) and concentrations were calculated by comparing peaks with corresponding calibration samples.

**Quantitative real-time PCR**

Medial PFC (mPFC) (+3.70 to +2.20mm from bregma), amygdala (-2.30 to -3.30mm) and ventral hippocampus (-4.80 to -6.04mm) [3] were isolated from brains stored at -80°C by micropunches (1.8mm of diameter). Total RNA was isolated from brain structures using TRIzol Reagent according to manufacturer instructions (ThermoFisher Scientific, Saint-Herblain, France). Reverse transcription was carried out on 2µg of total RNA using SuperScript III (Life technologies, Carlsbard, CA, USA). SYBR-green PCR kit (Eurogentec, Angers, France), specific primers pairs (supplementary table 1), and a previously validated quantity of cDNA were used for qPCR reactions in a Roche light Cycler 480 instrument. Relative mRNA levels were calculated using 2^-ΔΔCt^ method and represented as fold change.

**Golgi-cox staining**

Brains of Wistar and GK rats were rapidly immersed in the Golgi-Cox impregnation solution containing potassium dichromate, mercuric chloride and potassium chromate for 11 days before freezing in isopentane according to the manufacturer´s instructions (PK401 FD Rapid Golgi Stain KIT, Neurotechnology’s INC, Paris, France). mPFC (+3.72 to + 2.52 mm from bregma) were cut using a cryostat (Leica CM3050-5, Solms, Germany) to obtain coronal sections (200 µm) that were mounted in 2% gelatin-coated slides. Slides were incubated in Golgi-Cox staining solution (10 min, at room temperature, RT) with agitation, for being then immersed in a graded series of ethanol (50%, 75%, 95% and 100%) and xylene, and finally cover slipped using DPX mounting medium (EMS, Hatfield, PA). Throughout the process, light exposure was avoided.

A digital slide scanner (Nanozoomer, Hamamatsu Photonics, Massy, France) from the Bordeaux Imaging Center (CNRS-INSERM and Bordeaux University) was used to obtain images at 20x magnification that were further blind analyzed with ImageJ and the NeuronStudio (CNIC, Mount Sinai School of Medicine, NY, USA) software. The dendritic morphology and dendritic spine density analyses were conducted in the layer II/III pyramidal neurons of the mPFC (4-6 neurons per rat according to their antero-posteriority from bregma and mPFC subregion). Apical and basal dendritic trees of each selected neuron were separately reconstructed. If the number of neurons per animal was not sufficient, in order to draw the complete dendritic tree, either the basal or the apical tree from a specific neuron was analyzed. To do so, 20x magnification NDPI images (1 image / 4 µm in the z-axis) were converted into TIFF format using the NDPItools plugin of the ImageJ software. Then, selected images were opened in the NeuronStudio software and dendritic trees were manually reconstructed. Once drawn, Scholl analyses with ring radius increments of 20 µm were conducted to determine dendritic length for each reconstructed dendritic tree. ImageJ 1.52a software (http://imagej.nih.gov.gate2.inist.fr/ij/) was used to analyze density of dendritic spines in the same neurons. Dendritic spines were manually labelled on 80 µm segments and the density was expressed as the number of spines per 10 µm. For each selected neuron, 3 second-order and 2 third-order dendritic branches from the basal tree were studied. Regarding the apical tree 1 second-order, 2 third-order, 1 fourth-order and 2 fifth-order branches were examined. The experimenter remained blind to the treatment conditions throughout the analysis.

**Immunohistochemistry and quantification astrocytes activated neurons in the brain**

Rats received a lethal dose of pentobarbital to intracardially perfuse PBS 1X followed by paraformaldehyde 4% perfusion. Brains, previously immersed in paraformaldehyde 4% for 24h at 4°C and in 30% sucrose solution, were frozen in isopentane and stored at -80°C until processing. For astrocytes analysis, rats were sacrificed under basal conditions while the assessment of neuronal activity was explored in rats sacrificed 90 min after being exposure to a mild stressful stimulus (10 min in novel environment).

For all immunohistochemistry assessments, coronal brain sections (40µm), obtained in a cryostat (Leica CM3050S), were stored in cryoprotectant solution (glycerol and ethylene glycol diluted in PBS 1X) at -20°C in 10 consecutive series until immunolabeling procedure. One-in 10 series was used for either astrocytes or activated neurons immunolabelling using 3-5 sections of each brain region of interest per animal. Astrocytes were immunostained in mPFC (anterior-posterior, AP, mm from Bregma: +3.70 to +2.20) sections while activated neurons quantification was conducted in sections of mPFC, paraventricular nucleus (PVN) of the hypothalamus (AP: -0.92 to -2.12), amygdala (AP: -2.30 to -3.30) and ventral hippocampus (AP: -4.80 to -6.04).

Cryopreserved coronal brain sections were 1h-incubated with 1% bovine serum albumin (0.3% Triton-X100) followed by an overnight incubation (4°C) with 1:1000 dilution of rabbit anti- glial fibrillary acidic protein (GFAP) (Dako) or rabbit anti- early growth response 1 (EGR1) primary antibodies (Cell Signaling, Neuss, Germany). Sections were then incubated with 0.3% H_2_O_2_ (30 min room temperature, RT), followed by incubations with 1:2000 of biotinylated goat anti-rabbit secondary antibody incubation (2h at RT), and 1:1000 avidin-biotin complex (30-90 min at RT) (Vectastain ABC kit, Vector laboratories, Burlingame, CA). Immunostaining was revealed with diaminobenzidine via the nickel-enhanced glucose oxidase method.

Quantification of GFAP (astrocytes) and EGR1 (activated neurons) immunoreactive cells were quantified as follows: GFAP+ cells were quantified by design-unbiased stereology estimation using optical fractionators method [4] in each mPFC subregion (infralimbic, IL; prelimbic, PrL; and cingulate, Cg). Cells were visualized using 20x magnification in an Olympus BX51 microscope with a monitored Z and X-Y encoders linked to a software for stereological quantification (Mercator Pro software, Explora Nova, Universal laboratory imaging, La Rochelle, France) as previously described [5]. GFAP-immunolabelled cells were quantified in grids randomly placed in IL (size, 115x115 µm and spacing, 40x40 µm), (PrL and Cg (size: 175x175 µm and spacing: 100x100 µm) subregions. EGR1-immunolabelled cells quantification was conducted using Image J software. Regions of interest (ROI) were manually circumscribed using ROI tools command as follows: 3 subregions of the mPFC (IL, PrL and Cg), one region in PVN, 2 subregions of the amygdala (central nucleus, Ce; basolateral nucleus, BL), and 3 subregions of the ventral hippocampus (dentate gyrus, DG; CA1, CA3). EGR1-positive cells were automatically quantified in 8-bit thresholded images using the particle analysis function (15-100 µm²; circularity: 0.1-1).

**Anxiety-like and depressive like behaviors**

Wistar and GK male offspring behavior was assessed during the light phase. For analyses involving manual quantifications, experimenters remained blind to the experimental groups.

Anxiety-like behaviors:

*Light-dark test:*

The apparatus consisted in two connected compartments: a dark closed (31x31x45 cm high) and a light opened (illumination:100 lux, 45x31x45 cm high). Rats were placed in the dark compartment, and, after 10 min of exploration, behavior was video-recorded. The number of visits and the time spent in the light compartment were quantified using an ethological software (The observer, Noldus Information Technology, Wageningen, The Netherlands).

*Open-field test:*

Rats were placed for 10 min in the open-field arena (100×100 cm; illumination:100 lux). Distance and time spent in the center area were automatically assessed using a video tracking system (Viewpoint Behavior Technology, Lyon, France).

*Tunnel test:*

Rats were placed in a familiar environment (home cage with Sawdust and tube) for 5 min and behavior was video-recorded. The latency to enter in the tube, the number of entries and the time spent in the tube were quantified (The observer, Noldus Information Technology, Wageningen, The Netherlands).

Depressive-like behaviors:

*Social interaction*

Pairs of weight-matched rats from the same group (but unfamiliar) were placed in a neutral arena (40x40 cm) under dim light (16-30 lux) for 10 min. Time spent in social interaction (anogenital sniffing), the latency to social interaction and the duration of immobility were quantified (The observer, Noldus Information Technology, Wageningen, The Netherlands).

*Forced-swimming test (FST):*

Rats were placed in a cylinder (51 cm height x 21 cm diameter) filled to 29 cm with water (24+/-1 °C). A pretest of 15-min swimming session was performed and 24h later rats underwent a 5-min swimming test. At the end, rats were removed from the cylinder, dried and returned to their home cage. Duration of immobility, small movements (i.e. floating with small movements necessary to keep the head above water) and climbing defined as high activity (i.e. rigorous movements towards the cylinder walls resembling climbing behavior) were automatically quantified using a VideoTrack FST software (Viewpoint Behavior Technology, Lyon, France).

*Locomotor circadian rhythm*

Spontaneous locomotor activity throughout the light-dark cycle was assessed in rats (3 months old) for 48h using cages (18x30x18cm) equipped with photoelectric cells (Imetronic, Pessac, France). Rats were individually placed during 72h in the cage, the first 24h were used for habituation to the cage and were not recorded.

**Peripheral glucocorticoids normalization experiment**

Bilateral adrenalectomy or sham (used as controls) was conducted in anesthetized GK rats by a dorsal approach. To normalize the levels and circadian rhythm of glucocorticoids, low-dose corticosterone (25µg/mL in 0,9% saline) was provided in the drinking water as previously described [6] immediately after the surgery and during the experiment. To assess adrenalectomy efficacy, plasma corticosterone levels were measured in blood collected before and after a 15min-restraint stress using an ELISA kit [Corticosterone HS (High Sensitivity) kit (IDS®)]. Glycemia was also measured in the same time-points with ACCU-CHEK performa system (Roche Diagnostics, Mannheim, Germany). Insulin was measured in plasma at baseline with an ELISA kit (ALPCO, Salem, NH, USA). Corticosterone levels in PFC was measured as previously indicated. Behavioral assessments including light-dark, open-field, tunnel test, FST were performed between 25- and 90-days post-adrenalectomy as previously described. Social behavior was assessed by exposing experimental animals to a juvenile (postnatal day 40) for 10 min.

**Supplementary Table S1:** genes and primer pair sequences

| **Gene name** | **Coding for:** | | **Sequence 5´- 3´** | **cDNA(ng)** |
| --- | --- | --- | --- | --- |
| β2m  beta-2 microglobulin  (house keeping gene) | **β2M** | Forward: GCGTGGGAGGAGCATCAG  Reverse: TGTGCTGTAGGCCCAACAGA | | 2.5, 5, 25, or 40 |
| Ddit4  damage inducible transcript 4 | **REDD1** | Forward: CGCTCTTGTCCGCAATCTTC  Reverse: GGACGCTGGTTGATGAGGTT | | 5 |
| Fkbp5  FK506 binding protein 5 | **FKBP5** | Forward: GGCAGCCTCCCGAAAATT  Reverse: CTCCGGATAATGCCTGAATCTT | | 5 |
| Htr1a  5-hydroxytryptamine receptor 1A | **5HTR1A** | Forward: GCGTTGTTGGGTGCCATAAT  Reverse: CCGGATTGAGCAGGGAGTT | | 2.5 |
| Il1β  interleukin 1 beta | **IL1 β** | Forward: GACTTGGGCTGTCCAGATGAG  Reverse: TGAGTGACACTGCCTTCCTGAA | | 25 |
| Il6  interleukin 6 | **IL6** | Forward: ATATGTTCTCAGGGAGATCTTGGAA  Reverse: GTGCATCATCGCTGTTCATACA | | 25 |
| Nr3c1  nuclear receptor subfamily 3, group C, member 1 | **GR** | Forward: CTTTACGAAGTGTTTCTACTACTG  Reverse: TGACACCCAGAAGCCTCATCT | | 2.5 |
| Nr3c2  nuclear receptor subfamily 3, group C, member 2 | **MR** | Forward: GGGACCACCTCCCAAGCT  Reverse: ACCCCGTAATGACATCCT | | 2.5 |
| Sgk1  serum/glucocorticoid regulated kinase 1 | **SGK1** | Forward: GGGACAACGTCCACCTTCTG  Reverse: GGTCGTACGGCTGCTTATGG | | 5 |
| Socs3  suppressor of cytokine signaling | **SOCS3** | Forward: CACATGGCACAAGCACAAAAA  Reverse: GCTGGGCTAACTGGGAGCTA | | 5 |
| Tnfα  tumour necrosis factor alpha-like | **TNFα** | Forward: AGGCTGTCGCTACATCACTGAA  Reverse: TGACCCGTAGGGCGATTACA | | 25 |

Primers were designed with the Primer Express Software (Life Technologies, Applied Biosystems, PE Corporation, NY). Pre-designed primer pairs for *Htr2a* 5-hydroxytryptamine receptor 2A (**5HTR2A**), *Htr2c* 5-hydroxytryptamine receptor 2C (**5HTR2C**) *Slc6a4* solute carrier family 6 member 4 (**5HTT**) genes were provided by Applied Biosystems assay on-demand gene expression (California, USA) (40 ng of cDNA were use for the amplification)

**Supplementary Table S2:** sample size for qPCR

|  |  | **mPFC** | **AMYGDALA** | **HIPPOCAMPUS** |
| --- | --- | --- | --- | --- |
| ***Nr3c2*** | **Wistar** | n=8 | n=7 | n=7 |
|  | **GK** | n=8 | n=8 | n=8 |
| ***Nr3c1*** | **Wistar** | n=8 | n=7 | n=7 |
|  | **GK** | n=8 | n=8 | n=7 |
| ***Fkbp5*** | **Wistar** | n=8 | n=7 | n=7 |
|  | **GK** | n=8 | n=8 | n=6 |
| ***Ddit4*** | **Wistar** | n=8 | n=7 | n=8 |
|  | **GK** | n=8 | n=8 | n=6 |
| ***Sgk1*** | **Wistar** | n=8 | n=7 | n=8 |
|  | **GK** | n=8 | n=8 | n=5 |
| ***Il6*** | **Wistar** | n=7 | n=6 | n=6 |
|  | **GK** | n=8 | n=8 | n=7 |
| ***Socs3*** | **Wistar** | n=8 | n=7 | n=8 |
|  | **GK** | n=8 | n=8 | n=7 |
| ***Htr1a*** | **Wistar** | n=8 | n=6 | n=7 |
|  | **GK** | n=8 | n=8 | n=7 |
| ***Htr2a*** | **Wistar** | n=8 | n=7 | n=7 |
|  | **GK** | n=8 | n=7 | n=7 |
| ***Htr2c*** | **Wistar** | n=7 | n=7 | n=7 |
|  | **GK** | n=7 | n=7 | n=7 |
| ***Slc6a4*** | **Wistar** | n=8 | n=6 | n=7 |
|  | **GK** | n=8 | n=8 | n=7 |
